# T cells compete via reverse MHC class I signaling at the synapse with dendritic cells to secure Golgi recruitment for activation

**DOI:** 10.64898/2026.05.13.724773

**Authors:** Anthi Psoma, Elke M. Muntjewerff, Mara J.T. Nicolasen, Rjimke Bottema, Martin ter Beest, Rinse de Boer, Frans Bianchi, Natalia H. Revelo, Geert van den Bogaart

## Abstract

Interleukin-12 (IL-12) is a key immunostimulatory cytokine produced by dendritic cells (DCs) upon infection that plays a central role in the activation and differentiation of cytotoxic T cells. Notably, IL-12 is secreted in a highly polarized manner from late endosomes or lysosomes at the contact interface between a DC and a T cell, known as the immunological synapse. However, the signaling mechanisms initiating this spatially restricted cytokine release remain poorly defined. Here, using artificial T cell mimics, beads coated with T cell surface ligands, we show that antigen recognition induces recruitment of the DC’s microtubule organizing center (MTOC) and Golgi apparatus to the immunological synapse via reverse MHC class I signaling. Our results further indicate that the signaling pathways governing MTOC polarization are highly conserved in DCs, consistent with mechanisms described in other immune cell types. As newly synthesized IL-12 traffics through the Golgi, this intracellular reorientation facilitates localized release of IL-12 directly at the site of T cell engagement, likely enabling DCs to focus their stimulatory capacity on responsive T cells within the crowded environment of secondary lymphoid organs. Furthermore, by applying spatially patterned antibody arrays targeting MHC class I, we reveal that MTOC polarization is biased toward sites exhibiting the strongest reverse MHC class I signaling. This suggests that IL-12 is preferentially released at synapses formed with T cells displaying higher T cell receptor affinity, providing a mechanism for selection of the most potent T cell.

## Introduction

The activation of cytotoxic T lymphocytes (CTLs) requires the formation of an immunological synapse between a dendritic cell (DC) and a T cell. Here, peptide fragments derived from ingested or expressed proteins are presented in major histocompatibility complex class I molecules (MHC class I) on the surface of the DC to T cells in the secondary lymphatic organs (Angus & Griffiths, 2013; Friedl et al., 2005; Soares et al., 2013). This occurs via a process called repertoire scanning, where the DCs present peptides to a large number of T cells to identify T cell clones with cognate antigen-specific T cell receptors (TCRs) (Miller et al., 2004). Only T cells that express a cognate T cell receptor (TCR) recognizing the peptide fragment presented on MHC become activated and subsequently differentiate into effector CTLs. This activation and differentiation of T cells require not only the engagement of the TCR with the MHC-peptide complex, but also co-stimulation by membrane-bound receptors and soluble cytokines (Zehn et al., 2012). Particularly the cytokine interleukin-12 (IL-12) is important for activation of CTLs. IL-12 is a heterodimeric cytokine called IL-12p70, which is composed of the IL-12p35 and IL-12p40 subunits. IL-12 is produced by DCs that have been matured in response to inflammatory signals and is essential for host defence against infections with viruses and intracellular microbial pathogens and for control of malignancy (Liu et al., 2005; Trinchieri, 2003). IL-12 is important for activation of antigen-specific CTLs, particularly via induction of interferon (IFN)-γ expression (Rastogi et al., 2022).

Lymph nodes are very crowded organs that contain millions of T cells, but only a very small fraction of these T cells carries a TCR recognizing a particular epitope, which can be as low as 1 in a million (Jenkins & Moon, 2012). Upon reaching the lymph nodes, naive T cells survey DCs for cognate peptide-MHC complexes and co-stimulatory signals through multiple transient, serial contacts known as kinapses, before forming stable, long-lasting interactions called synapses. These stable immune synapses are thought to be essential for complete T cell differentiation (Sapoznikov et al., 2023). However, prolonged DC-T cell interactions are not essential for T cell activation; activation can occur during the initial phase of transient contacts (Verboogen et al., 2016). These interactions can be summarised in three major phases: In phase 1 (up to 8 hours after entry into the lymph node), motile naïve T cells actively scan multiple DCs in search of their cognate antigen. During phase 2 (8–24 hours), stable, sessile interactions form between antigen-specific T cells and DCs, often lasting several hours. In phase 3 (24–48 hours), activated T cells detach from DCs, regain motility, and begin to proliferate (Chudnovskiy et al., 2019).

In order to avoid bystander activation of nearby T cells without a cognate TCR, a large fraction of IL-12 is released in a highly polarized fashion at the immunological synapse with antigen-specific T cells (Chiaruttini et al., 2016; Pulecio et al., 2010). An important event that underlies the recruitment of cytokines to the immunological synapse between DCs and T cells is the general reorganization of the cell’s cytoskeleton (Benvenuti, 2016). During this process, the microtubule organizing center (MTOC) migrates towards the immunological synapse in both the DC and T cell due to clearance of actin at the center of the synapse, leading to re-arrangement of the microtubular network. This reorientation of the MTOC is paired with the translocation of the Golgi apparatus and secretory machinery towards the immunological synapse (Dustin et al., 2010), enabling the localized release of cytokines such as IL-12 and IL-6 by antigen presenting cells (Dustin et al., 2010; Verboogen et al., 2016). Following its assembly in the endoplasmic reticulum (ER) and Golgi apparatus, vesicles with newly synthesized IL-12 gather around the MTOC and the associated Golgi apparatus and are transported towards the immunological synapse in a process that depends on the Rho-GTPase Cdc42, a well described polarization molecule, and on remodeling of the microtubular cytoskeleton (Borg et al., 2004; Pulecio et al., 2010). The final secretion at the immunological synapse occurs from vesicles of late endosomal or lysosomal nature by the action of the SNARE protein VAMP7 (Chiaruttini et al., 2016).

So far, MTOC polarization towards the synapse has been described in multiple immune cell types, including DCs, helper T cells, CTLs, B cells and natural killer (NK) cells. For most of these cell types the initial signaling molecules that trigger polarization are known. For example, MTOC polarization is triggered by the B cell receptor for B cells (Wang et al., 2017) and CD28, NKG2D, NKp30 and CD94 for NK cells (Chen et al., 2007). MTOC polarization is best understood for T cells, where TCR engagement initiates a protein kinase C (PKC) dependent signaling cascade that induces cytoskeletal reorganization by dynein-dependent trafficking of the MTOC towards the antigen presenting cell (APC) (Huse, 2012). However, for DCs, little is known about the signaling cascade involved in their MTOC polarization. So far it has been shown that this process is antigen dependent, it only takes place when the DC is activated and that it involves the conserved polarization GTPase Cdc42 (Pulecio et al., 2010). What initial molecule(s) triggers MTOC polarization and results in the stable synapse formation in DCs remains to be elucidated. In this study, we show that the polarized release of IL-12 results from the migration of the MTOC together with the Golgi apparatus towards the immunological synapse. Moreover, we show that MTOC polarization is mediated by reverse MHC class I signaling, and its recruitment is proportional to the strength of this reverse signaling.

## Results

### MTOC and Golgi polarization in DCs at synapses with T cells

To visualize the polarization of the MTOC within the DCs towards the immunological synapse with T cells, we used a well-established synapse model based on murine OT-I T cells that recognize the peptide SIINFEKL, derived from the model antigen ovalbumin (OVA), when presented in the context of MHC class I (H-2Kb) (Fig. 1A). To induce synapse formation, murine Flt3L-differentiated bone-marrow derived DCs (BMDCs) were isolated from wild-type mice, activated with the Toll-like receptor ligands lipopolysaccharide (LPS) and CpG_B 1826 and loaded with SIINFEKL. As control treatments for antigen processing, BMDCs were also loaded with the full OVA protein or OVA-antibody immune complexes. These cells were then incubated with OT-I T cells to allow for synapse formation. The cells were then immunostained for α-tubulin, γ-tubulin and GM130 to identify the microtubular network, the MTOC, and the Golgi apparatus, respectively (Fig. 1B). As done previously (Pulecio et al., 2010), the MTOC polarization index was calculated by measuring the distance between the MTOC and the DC-T cell interface divided by the diameter of the DC perpendicular to the axis of the synapse (Fig. 1C). DCs with a polarization index equal to or below 0.3 were considered polarized and displayed as percentage of polarized cells (Fig. 1D-E), similar to previous analysis (Pulecio et al., 2010). In line with this previous research (Pulecio et al., 2010), MTOC polarization to the immunological synapse was more predominant in DCs that were both activated with TLR ligands and loaded with antigen. TLR activated DCs that were not loaded with antigen did not show polarization (Fig. 1D-E). To confirm synapse formation and the positioning of the MTOC and Golgi at the immunological synapse, we performed transmission electron microscopy on interacting murine OT-I T cells and BMDCs (Fig. 1F). We observed a smooth and tight cellular interface between the cells. Additionally, we observed that the MTOC and Golgi apparatus located close to the DC-T cell synapse in the DC, indicating the effector stage.

**Figure 1.**
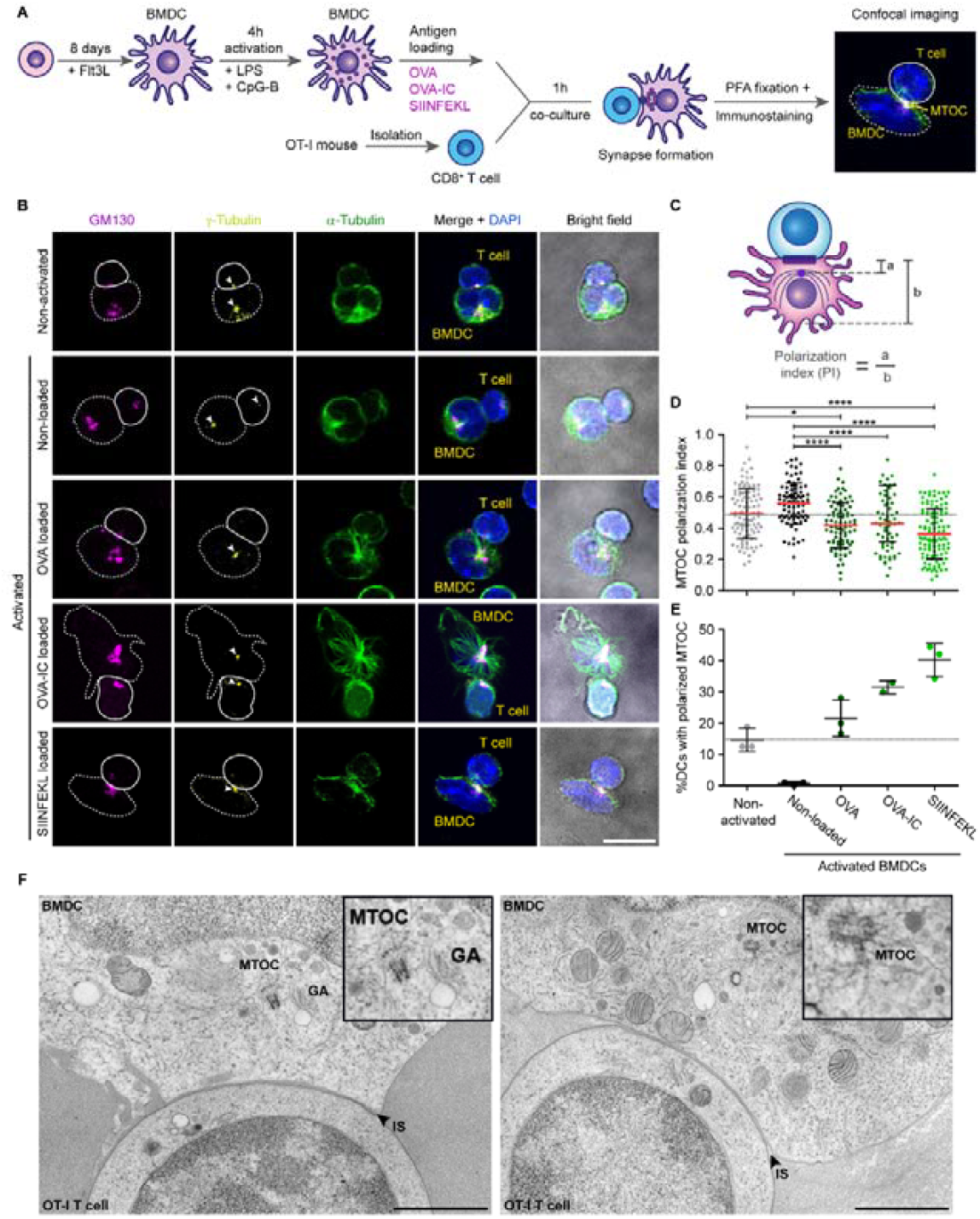
MTOC and Golgi polarisation in DCs at immunological synapses with T cells. **(A)** Scheme of synapse formation between murine Flt3L-differentiated bone marrow-derived DCs (BMDC) and OT-I T cells. LPS: lipopolysaccharide; CpG-B: oligodeoxynucleotide; OVA: ovalbumin; IC: immune complex; SIINFEKL: ovalbumin (257-264) peptide epitope; PFA: paraformaldehyde. **(B)** Representative confocal microscopy images of the BMDC-T cell synapses immunostained for cis-Golgi marker GM-130 (magenta), γ-tubulin (yellow), DAPI (blue) and α-tubulin (green). White arrowheads: MTOC polarization. Scale bar, 10 µm. **(C)** Scheme of quantification of MTOC polarization: the distance between the MTOC and the DC-T cell interface (a) was divided by the diameter of the DC (b). **(D)** Quantification of MTOC polarization with unstimulated BMDCs or TLR ligand-stimulated BMDCs unloaded or loaded with OVA, OVA-Immunocomplexes or SIINFEKL. For each of the 3 mouse cultures, 30-35 cells were analysed. Mean ± standard error of the mean, Kruskal-Wallis test, Dunn’s multiple comparison **P<0.01, ****P<0.0001 **(E)** Same as panel D, but now displayed as percentage DCs with polarized MTOC (defined as polarization index < 0.3) for individual mice. **(F)** Electron micrographs of two different immunological synapses between T cell and BMDC, showing MTOC, Golgi apparatus (GA), and immunological synapse (IS); Black boxes indicate magnified areas. Scale bars, 1 µm.

### Reverse MHC class I signaling drives MTOC polarisation in murine DCs

Synapse formation results in MTOC polarization in DCs, however the signals leading to this are unknown. Based on the receptors triggering MTOC polarization in B, T and NK cells (Chen et al., 2007; Huse, 2012; Wang et al., 2017), we selected the following candidates on the DC membrane that might potentially trigger MTOC polarization: Antigen presentation receptors (MHC class I and class II) (Gur et al., 1990; Tsai & Reed, 2014; Wagner et al., 1994), costimulatory molecules CD80 and CD86 (Sedwick et al., 1999), and adhesion molecules LFA-I and ICAM-I (Pulecio et al., 2010; Sedwick et al., 1999). To identify the initial molecule(s) leading to MTOC polarization, we developed a model where mouse Flt3L-differentiated BMDCs were stimulated with antibody-coated beads or protein domain-coated beads (Fig. 2A). Non-activated BMDCs and BMDCs stimulated with naked beads or isotype control antibody-coated beads were included as negative control conditions. To visualize the MTOC polarization in our bead synapse model, cells were again immunostained for the microtubular network, the Golgi apparatus and the MTOC (Fig. 2B). Quantification of the polarization index, using the contact site with the bead as reference, revealed that mouse BMDCs only polarized when conjugated to beads coated with an antibody against MHC class I and not with antibodies recognizing MHC class II and ICAM-1 or protein domains of CD28 (ligand of CD80 and CD86) or LFA-I (Fig. 2C).

**Figure 2.**
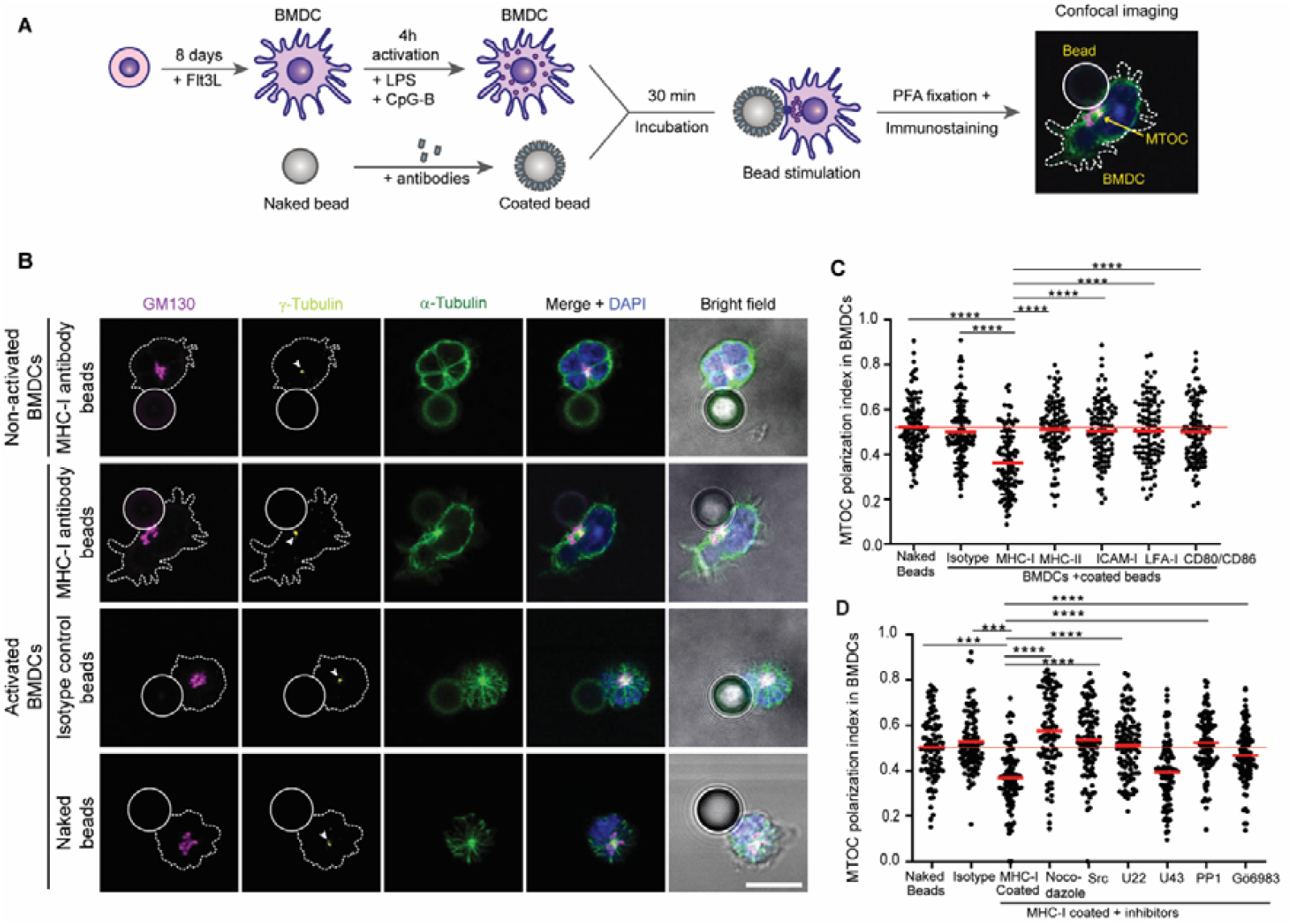
Reverse MHC class I signaling drives MTOC polarisation in murine DCs. **(A)** Scheme of MTOC polarization by antibody- or protein domain-coated bead stimulation of murine Flt3L-differentiated bone marrow-derived DCs (BMDCs). LPS: lipopolysaccharide; CpG-B: oligodeoxynucleotide; PFA: paraformaldehyde. **(B)** Representative immunofluorescence images of the BMDC-bead conjugates. Cells were immunostained for cis-Golgi marker GM-130 (magenta), γ-tubulin (yellow), DAPI (blue) and α-tubulin (green). Beads were coated with isotype or antibody against MHC class I. Straight outline: T cell; dotted outline BMDC; white arrowhead: MTOC polarization. Scale bar, 10 µm. **(C)** Quantification of MTOC polarization in BMDCs incubated with beads coated with antibodies against MHC class I, MHC class II, or ICAM-1, or with LFA-I or CD28 protein domains. One-way ANOVA Kruskal-Wallis test with Dunn’s Multiple Comparisons. **(D)** Quantification of MTOC polarization in BMDCs incubated with beads coated with antibodies against MHC class I treated with nocodazole inhibitor targeting the microtubules (Mt), the Src-I1 and PP1 inhibitors targeting tyrosine kinase Src, and the phospholipase C inhibitor (U73122; U22) and its negative control compound (U73343; U43) and the PKC inhibitor Gö6983. One-way ANOVA with Dunnett’s multiple comparison, Mean ± standard error of the mean (**P<0.01; ****P<0.0001).

Reverse MHC class I signaling is a process in which engagement of MHC class I molecules on the cell surface transmits signals into the same cell (Muntjewerff et al., 2020). This phenomenon has been described in immune cells such as macrophages, DCs, NK cells, B cells, and T cells, where it has been linked to modulation of immune responses, as well as in non-immune cells like endothelial and epithelial cells. Signaling through MHC class I is thought to be mediated by phosphorylation of a preserved tyrosine residue within the short cytoplasmic tail of the α heavy chain, which has been suggested to undergo phosphorylation by Src kinase in mouse and human cells (Guild et al., 1983; Santos et al., 2004; Xu et al., 2012). Although the signaling cascade ensuing MHC class I phosphorylation has been widely studied in non-immune cells, and to some extent in T and B lymphocytes, less is known in immune myeloid cells (Muntjewerff et al., 2020). In mouse macrophages, a crosstalk between TLR and MHC class I signaling was described, where TLR stimulation induces Src phosphorylation of MHC class I Y320 residue, which in turn recruits Fps kinase and leads to a signaling cascade that downregulates TLR-induced signals, including less activation of MAPK, NF-κB and IRF3 pathways (Xu et al., 2012). To assess if Src phosphorylation of MHC class I could be involved in MTOC polarization in DCs, we treated BMDCs with two Src inhibitors before stimulation with beads coated with anti-MHC class I antibodies (Src-I1 and PP1). Further studies demonstrated that the lipid second messenger diacylglycerol (DAG) accumulates as well in a polarized manner at the immunological synapse (Spitaler et al., 2006). In T cells, DAG is produced downstream of TCR activation via phospholipase C-γ (PLCγ), and inhibition of PLCγ has been shown to completely abrogate MTOC polarization in this cell type (Quann et al., 2009). These findings identified DAG signaling as a central regulator of cytoskeletal polarization. Based on this, we investigated whether the same pathway is conserved in DCs using the PLC inhibitor (U73122) and its negative control compound (U73343). The discovery that DAG regulates MTOC polarization also implicated DAG-responsive proteins containing C1 domains, particularly members of the protein kinase C (PKC) family (Huse, 2012). Subsequent work revealed a coordinated recruitment of PKCs at the immunological synapse, where PKC-ε and PKC-η accumulated rapidly over a broad membrane region, followed by the later recruitment of PKC-θ to a more confined zone. In addition, pharmacological inhibition of PKCs using Gö6983 strongly impaired MTOC polarization, while knockdown of PKC-θ resulted in pronounced reorientation defects (Quann et al., 2011). Building on these findings, we employed Gö6983 to investigate whether PKC signaling similarly regulates MTOC polarization in DCs. We also used nocodazole as a positive control of MTOC polarization disruption.

BMDCs incubated for two hours with the inhibitors were then stimulated with beads coated with antibody against MHC class I, fixed and assessed using immunofluorescence microscopy. As before, MHC class I-coated beads induced significant MTOC polarization when compared to isotype antibody-coated and uncoated beads (Fig. 2D). Inhibition of microtubule polymerization and Src activity both blocked this polarization, suggesting that Src could mediate MTOC polarization through phosphorylation of MHC class I in BMDCs. Furthermore, phospholipase C inhibitor and the PKC inhibitor inhibited MTOC polarization as well. Our results suggest that the signaling pathways controlling MTOC polarization are highly conserved across diverse immune cell types. Notably, our findings further demonstrate that MHC class I reverse signaling participates in MTOC reorientation in DCs.

### Reverse MHC class I signaling drives MTOC polarisation in human DCs

The tyrosine in the cytosolic region of MHC class I is present in all major MHC class I forms in mouse (H-2K, H-2D, H-2L) but only in two out of the three human forms (only in HLA-A and HLA-B, absent in HLA-C) (Fig. 3A). We therefore tested whether reverse MHC class I signaling also drives MTOC polarization in human DCs. We generated DCs from blood-isolated monocytes (moDCs), incubated them with beads coated with either an antibody against MHC class I or an isotype control and immunostained them for α-tubulin (Supplementary Fig. 1A) or the centrosome protein pericentrin (Fig. 3B-C). Quantification of the MTOC polarization index showed that, similar to mouse BMDCs, human DCs polarize when stimulated with beads coated with MHC class I antibody but not when stimulated with naked or isotype antibody-coated beads (Fig. 3D, Supplementary Fig. 1B-C). These data further suggest that MTOC polarization during synapse formation in mouse and human DCs is induced by reverse MHC class I signaling. Confirming our results with muring BMDCs, the microtubule polymerization inhibitors colchicine and nocodazole blocked MTOC migration to the synapse (Fig. 3D). We also tested MTOC polarization in a mixed lymphocyte reaction (MLR) as a model of immune synapse, by coincubating moDCs and peripheral blood lymphocytes (PBLs) from different donors. There, nocodazole and colchicine also could block MTOC polarization to the immunological synapse in moDCs (Fig. 3E).

**Figure 3.**
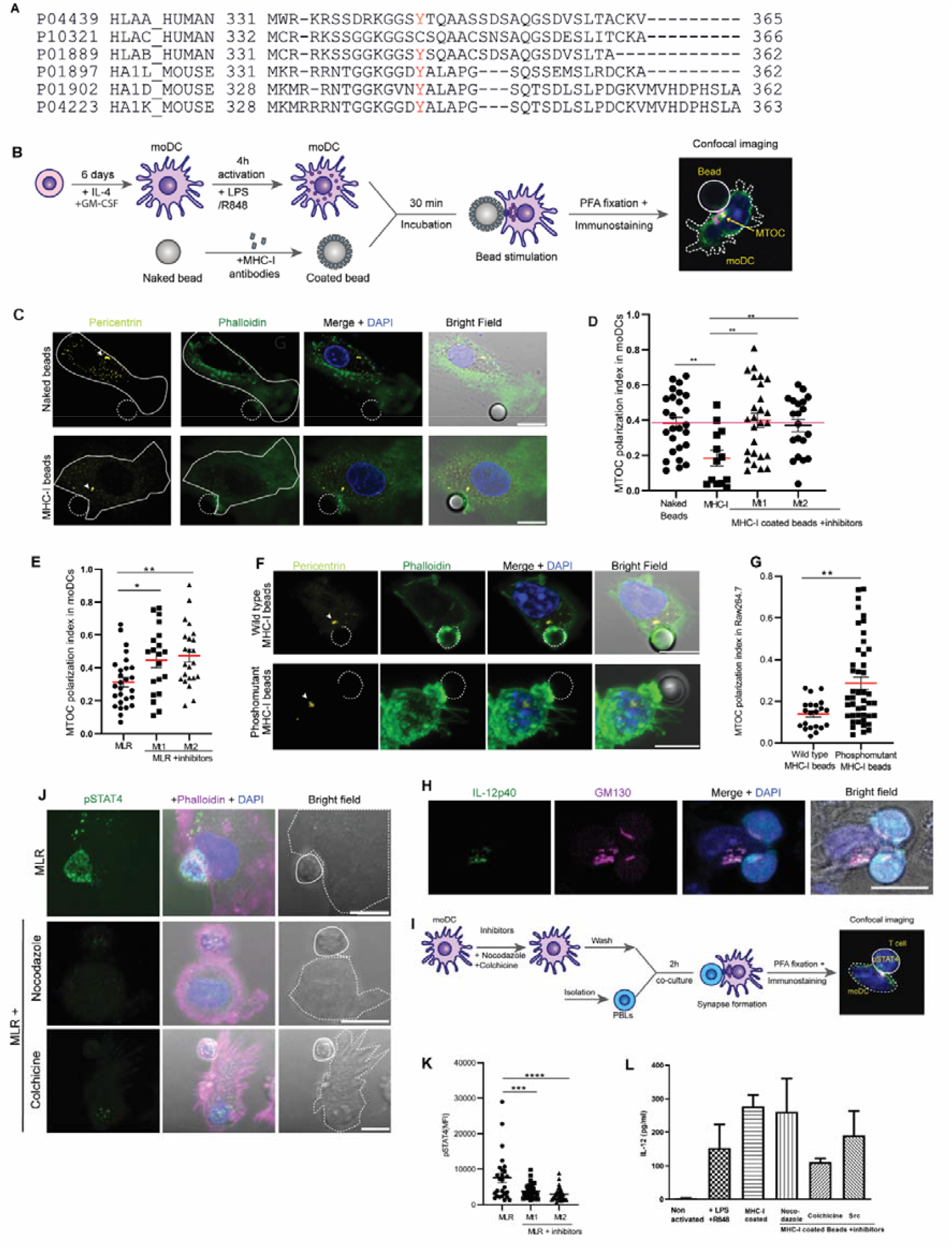
Reverse MHC class I signaling drives MTOC polarisation in human DCs. **(A)** Alignment of the C-terminal cytosolic tails of the main human and murine MHC class I forms. The Uniprot accession numbers are shown in the figure. The tyrosine is conserved in most, but not all, MHC class I forms. **(B)** Scheme of synapse formation between IL-4/GM-CSF-differentiated human monocyte derived DCs (moDC) and beads coated with antibodies recognizing MHC class I. LPS: lipopolysaccharide; R848: Resiquimod; IC: immune complex; PFA: paraformaldehyde. **(C)** Representative confocal microscopy images of the moDC-bead conjugates. Cells were immunostained for DAPI (blue), pericentrin (yellow), and phalloidin (green). Beads were naked or coated with antibody against MHC class I. Solid outline: moDC; dashed outline: bead; white arrowhead: MTOC polarization. Scale bars, 10 µm. **(D)** Quantification of the MTOC polarization in moDCs incubated with beads coated with antibodies against MHC class I and treated with the microtubule inhibitors nocodazole (Mt1) or colchicine (Mt2). One-way ANOVA with Dunnett’s multiple comparison. **(E)** Quantification of the MTOC polarization in moDCs incubated with PBLs treated with nocodazole or colchicine. One-way ANOVA with Dunnett’s multiple comparison. **(F)** Representative confocal microscopy images of the RAW 264.7 macrophages heterologously expressing human HLA-A*02:01 and incubated with bead coated with antibodies against human (but not mouse) MHC class I or with naked beads. Cells were immunostained for DAPI (blue), pericentrin (yellow), and phalloidin (green). Solid outline:bead; white arrowhead: MTOC polarization. Scale bars, 10 µm. **(G)** Quantification of the MTOC polarization toward beads coated with antibodies against human (but not mouse) MHC class I in RAW 264.7 cells expressing either wild type or phosphodead (Y344F) human HLA-A*02:01. Unpaired t test. **(H)** Representative confocal microscopy images of the BMDC-T cell synapses immunostained for cis-Golgi marker GM-130 (magenta), and IL-12p40 (green) and DAPI (blue). Scale bar, 10 µm. **(I)** Scheme of synapse formation between human derived DC (moDC) and T cells in a MLR reaction. PBLs: Peripheral blood lymphocytes; PFA: paraformaldehyde. **(J)** Representative confocal microscopy images of the MLR reaction. Cells were immunostained for DAPI (blue), pSTAT4 (green), and phalloidin (magenta). Scale bars, 10 µm. **(K)** Quantification of pSTAT4 mean fluorescence intensity (MFI) in PBLs incubated with moDCs treated with nocodazole (Mt1), colchicine (Mt2). One-way ANOVA with Dunnett’s multiple comparison. **(L)** Total IL12p70 production in moDCs incubated without LPS+ R848 and without beads, with LPS + R848 and without beads or with beads coated with antibodies against MHC class I and treated without or with the microtubule inhibitor nocodazole (Mt1), the colchicine inhibitor (Mt2) or Src kinase inhibitor (*P<0.05; **P<0.01; ***P<0.001; ****P<0.0001).

In order to confirm the role of phosphorylation of the conserved cytosolic tyrosine in reverse MHC class I signaling (Guild et al., 1983; Santos et al., 2004), we tested whether mutation of this amino acid would inhibit MTOC polarization to the immunological synapse. Because both murine BMDCs and human moDCs are difficult to transfect with sufficiently high efficiency, we used the murine macrophage-like cell line RAW 264.7 for this purpose. We transfected these cells with a plasmid coding for Myc-tagged human wild type HLA-A*02:01 (McShan et al., 2019) and a phosphomutant with tyrosine 344 replaced by a phenylalanine (Y344F). We incubated the cells with beads coated with an antibody recognizing human MHC class I or uncoated beads and immunolabeled the cells for the Myc-tag (Supplementary Fig. 2A) and the MTOC protein pericentrin (Fig. 3F). As the antibody only binds to human and not murine MHC class I, these experiments allowed us to test the effect of the tyrosine without interference from endogenous MHC class I forms. Whereas antibody binding to wildtype HLA-A*02:01 resulted in polarization of the MTOC towards the bead, this polarization was less for cells expressing the tyrosine mutant (Fig. 3G, Supplementary Fig. 2B-C), confirming that reverse MHC class I signaling is mediated by tyrosine phosphorylation.

### Golgi polarisation drives polarized release of IL-12

The p35 and p40 subunits of IL-12 are synthesized separately in the ER, where they fold, assemble, and form the biologically active IL-12p70 heterodimer (Jalah et al., 2013). From the ER, IL-12 traffics through the Golgi apparatus, where it undergoes glycosylation (Carra et al., 2000). Rather than being released directly from the Golgi, IL-12 is then sorted into late endosomal or lysosome-related secretory vesicles that are marked by the protein LAMP1 (Chiaruttini et al., 2016). These vesicles move along microtubules toward the plasma membrane where the SNARE protein VAMP7 mediates their fusion with the membrane to release IL-12 into the intercellular space between the DC and the T cell (Chiaruttini et al., 2016). As our data show that the Golgi apparatus migrates toward the immunological synapse within the DCs, this suggests that the polarised release of IL-12 could be the facilitated through its Golgi trafficking. To assess this, human monocyte-derived DCs were immunostained for IL-12p40. Indeed, we observed clear overlap of IL-12p40 with Golgi marker GM130. Thus, the formation of an immunological synapse with T cells results in recruitment of the Golgi apparatus to the synapse, which in turn could result in the polarised release of IL-12 (Fig. 3H).

IL-12 signals via phosphorylation of the transcription factor STAT4. We therefore performed a functional test to assess whether disruption of MTOC polarization in DCs with the microtubule inhibitors used above also disrupt phosphorylation of STAT4 in PBLs as a result of less IL-12 secretion at the synapse from moDCs. For this purpose, we initially tested pSTAT4 phosphorylation in human PBLs upon stimulation with IL-12. We incubated human derived PBLs with 500 ng/ml human IL-12 for one hour, and assessed STAT4 phosphorylation with confocal imaging (Supplementary Fig. 2D). We then tested pSTAT4 levels in a MLR reaction. First, moDCs were incubated with nocodazole or colchicine, washed to remove the inhibitors, and then co-cultured with PBLs (Fig. 3I). When compared to a negative control with no inhibitors, the results show that disruption of MTOC polarization leads to reduced STAT4 signaling in PBLs (Fig. 3J-K). This reduction is due to reduced polarized IL-12 release, because total IL-12 secretion was not affected by the microtubule inhibitors (Fig. 3L).

### MTOC polarisation at the site of strongest reverse MHC class I signaling

Live imaging of intact lymph nodes with two-photon microscopy has shown that once DCs enter a lymph node, they can transiently interact with over 500 T cells per hour during the initial phase of T cell activation (Bousso & Robey, 2003; Miller et al., 2004). Later during the T cell activation process, antigen-loaded DCs can form stable synapses with multiple peptide-specific T cells, engaging with more than ten simultaneously in interactions that can last for hours (Bousso & Robey, 2003). This results in synaptic-like cell-cell communication clusters where one DC interacts with many T cells, called rosettes, which are commonly observed in lymph nodes (Chudnovskiy et al., 2019). Also, in our fluorescence and electron microscopy experiments, we occasionally observed multiple T cells interacting with the same DC (Fig. 4A-B). However, this raises the question towards which of the T cells the MTOC of the DC would polarize, thus how a single MTOC manages competing targets.

**Figure 4.**
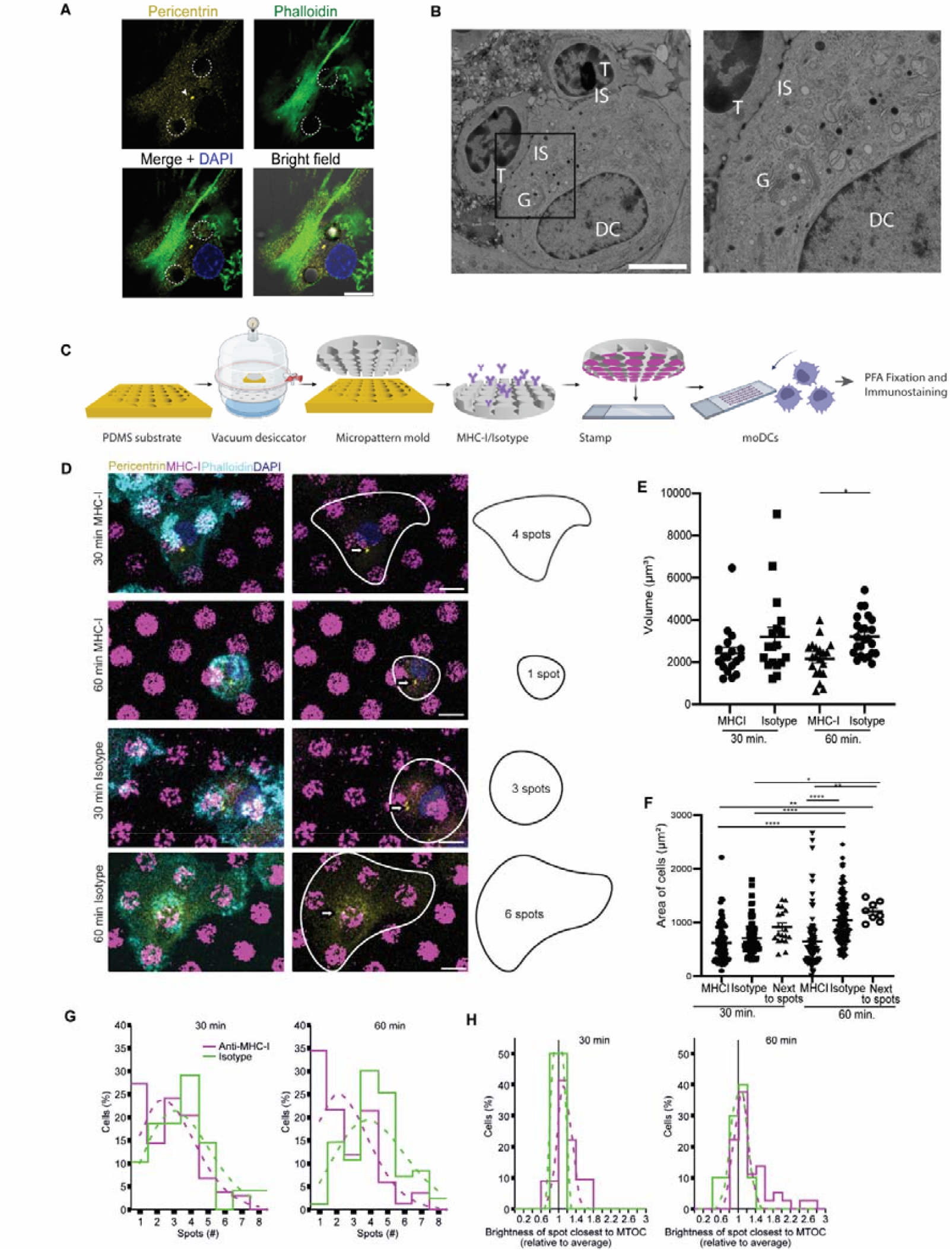
MTOC polarisation to the site of strongest reverse MHC class I signaling. **(A)** Confocal microscopy image showing a single human monocyte-derived DC (moDC) interacting with beads coated with antibodies against MHC class I simultaneously. Dotted outline: MHC class I bead. Scale bar, 10 µm. **(B)** Electron micrographs of immunological synapse between two murine OT-I T cells and a single bone-marrow derived DC presenting the cognate ovalbumin antigen. G: Golgi; IS: immunological synapse. Square: magnified area. Scale bar, 5 µm. **(C)** Scheme of PDMS stamps for generation on antibody patterns. The stamps were covered with MHC class I or isotype antibodies. MoDCs were cultured on the patterns. **(D)** MoDCs were seeded on patterns of MHC class I or isotype antibodies (purple) for the indicated times. Cells were immunostained for phalloidin (cyan), pericentrin (yellow), and DAPI (blue). Solid outline: moDC surface area. **(E)** Analysis of the volume of the cells seeded in MHC class I or control stamps at the indicated time points. For each donor, 6-8 cells were analysed. Ordinary one-way ANOVA, Tukey’s multiple comparison multiple comparisons), n = 3 donors. **(F)** Analysis of the projected surface area of the cells seeded on the MHC class I or isotype control antibody patterns for the indicated times. For each donor, 28-40 cells were analysed. One-way ANOVA with Tukey’s multiple comparison. (*P<0.05, **P<0.01, ***P<0.001), n = 3 donors. **(G)** Quantification of the number of spots covered by individual moDCs. Data were fitted with Poisson distributions (dashed lines). **(H)** Quantification of the fluorescent intensities of the spot closest to the MTOC divided over the average fluorescence intensities of all spots covered by that cell. Data were fitted with Gaussian distributions (dashed lines).

To obtain a quantitative understanding of MTOC polarization in situations where a DC is engaged in multiple synapses simultaneously, we used a technique called microcontact printing (Zuidscherwoude et al., 2017). This technique is a form of soft lithography that uses the relief patterns on a polydimethylsiloxane (PDMS) silicone elastomer stamp to print molecules of interest in two dimensional patterns on a microscopy support, such as a glass coverslip. This technique allows the assessment of cellular responses to spatially-restricted stimuli. We used a suspension of antibodies directed against MHC class I as stamping medium to create a grid of spots with a period of 16 µm and a diameter 8 µm on which human moDCs were cultured. This thus allows the spatially restricted response of moDCs to be assayed by creating artificial rosettes, as they encounter multiple areas coated with antibodies triggering reverse MHC class I signaling (Fig. 4C). An advantage of this approach is that all the printed areas are positioned in the same focal plane, thus allowing for quantitative microscopy approaches. As a negative control we generated patterns with an isotype antibody. Following the seeding, we incubated the cells for 30 and 60 minutes. After these time points the cells were fixed and immunostained for pericentrin (Fig. 4D).

We first determined the morphological characteristics of the cells, because DCs are well known to change their shape during immunological synapse formation, as they round up and cover less surface area (Verboogen et al., 2016). Although we observed only small differences in the surface area (2D-projected sizes) and volume (calculated from confocal microscopy stacks) of the DCs at 30 min after seeding, after 60 min we noticed that MHC class I engagement resulted in a substantially lower surface area compared to the isotype control antibody (Fig. 4E-F). Moreover, the spreading of the cells was also reduced compared to the cells that adhered adjacent to the patterns on the same cover slip (Fig. 4F). Opposite to the cells on the patterns, the cell area of the cells adjacent to the patterns increased over time (Fig. 4F). These findings were corroborated by the number of antibody spots covered by each cell, because after 60 minutes of incubation the cells that were seeded on the MHC class I antibody spots covered less spots than the cells that were seeded on the isotype antibody control spots (Fig. 4G).

Moreover, as we generated patterns with fluorescently-labeled antibodies, the fluorescence intensity of each spot is proportional to the number of antibodies and hence the potency to induce reverse MHC class I signaling (i.e., a high fluorescence intensity correlates with high reverse MHC class I signaling). Therefore, we measured the integrated fluorescence intensity of each spot that was covered by a cell (Fig. 4D). We then divided the fluorescence intensity of the spot that was located closest to the MTOC over the average intensity of all the other spots that were covered by the same cell. Values above one thus indicate that the MTOC is located at brighter spots with higher amounts of anti-MHC class I antibody. We noticed that after 1 hour incubation, the MTOC was most often located closer to the brighter MHC class I spots whereas this was not the case for the isotype control spots (Fig. 4H). Thus, we conclude that when DCs engage in multiple immunological synapses simultaneously, the MTOC polarizes to the site of strongest reverse MHC class I signaling.

## Discussion

DCs are the central orchestrators of the adaptive immune response, serving as the primary activators of naive T cells (Chaplin, 2010; Norcross, 1984). In their immature state, DCs continuously survey peripheral tissues for pathogens or malignant cells, capturing them through phagocytosis and endocytosis. Upon recognition of danger signals via pattern recognition receptors (PRRs) or inflammatory cytokines, DCs undergo activation and maturation. This process drives a phenotypic switch that enables their migration to secondary lymphoid organs, such as lymph nodes and spleen, where they engage with and activate T cells. The initiation of T cell activation requires three distinct but coordinated signals, called signals 1–3. Signal 1 is the antigen recognition through T cell receptor binding to the cognate peptide-MHC complexes. Signal 2 is co-stimulatory signaling via CD80/CD86 interactions with CD28. Signal 3 is cytokine-mediated instruction that potentiates activation and directs T cell differentiation into specific effector subsets (e.g., CTL, Th1, Th2, Th17, or Treg). Through this three-signal system, DCs integrate innate sensing cues to tailor adaptive immune responses appropriately to the nature of the encountered pathogen.

Within the complex environment of the lymph node, DCs must efficiently activate antigen-specific T cells among a vast repertoire. This is achieved through the polarized release of signaling molecules toward the immunological synapse (Verboogen et al., 2016). Although less extensively studied than T cell polarization, DC-side trafficking and polarization promote efficient antigen presentation and signal delivery. During synapse formation, DCs locally traffic MHC class I and II molecules, as well as co-stimulatory molecules such as CD40, to the contact site, allowing T cells to detect even minimal numbers of cognate MHC-peptide complexes (Bertho et al., 2003; Boes et al., 2002, 2003; Compeer et al., 2014; Foster et al., 2012). Additionally, IL-12 secretion at the synapse enhances Th1 polarization, promotes cytotoxic activity of CD8□ T cells, and induces IFN-γ production (Pulecio et al., 2010). MHC class II trafficking to the synapse follows a well-defined pathway: after synthesis in the ER and Golgi, MHC II associates with the invariant chain (Li) and transits to the MHC class II compartment (MIIC) for peptide loading (Berger & Roche, 2009; Neefjes et al., 2011; Rocha & Neefjes, 2008). Upon maturation, pathogen-induced signaling (e.g., via TLR4) drives MIIC tubulation toward the plasma membrane, a process regulated by Rab7 and Arl8b (Mrakovic et al., 2012). These tubular structures orient toward the immunological synapse under the influence of TCR-MHC and LFA-1-ICAM interactions, ensuring precise spatial delivery of MHC class II (Bertho et al., 2003).

In contrast, the mechanisms guiding polarized cytokine trafficking and secretion at the immunological synapse remain less clearly defined. Our study identifies reverse MHC class I signaling as a key initiator of the cytoskeletal remodeling required for the polarization of the MTOC and the associated Golgi apparatus to the immunological synapse. Reverse MHC class I signaling has been increasingly recognized as a pivotal pathway modulating both cytoskeletal organization and innate immune activation (Muntjewerff et al., 2020). In DCs, MHC class I engagement during antigen-specific synapse formation with CD8^+^ T cells induces extensive remodeling of endosomal recycling compartments (ERCs), transitioning from vesicular to tubular structures that facilitate targeted MHC class I transport to the synapse (Compeer et al., 2014). Similar remodeling can be elicited experimentally through antibody-mediated ligation of MHC class I or ICAM-1, particularly in LPS-activated DCs, indicating that MHC class I functions as a signaling hub coordinating microtubule and vesicular dynamics (Compeer et al., 2014).

Reverse MHC class I signaling has also been described in macrophages. Early work demonstrated that antibody-mediated targeting of MHC class I induces reactive oxygen species (ROS) production in macrophages through a process known as bipolar bridging, where antibodies engage both MHC class I and Fc receptors on the same cell membrane (Benichou & Voisin, 1987). Although this mechanism represents a collaboration between MHC class I and Fc receptors rather than pure reverse MHC class I signaling, later studies elucidated a defined inhibitory pathway mediated by MHC class I itself. Specifically, phosphorylation of the tyrosine residue in the cytosolic region of MHC class I recruits the tyrosine kinase Fps, which subsequently activates the phosphatase SHP-2 (Xu et al., 2012). SHP-2 interacts with TRAF6 to reduce its ubiquitination and downstream NF-κB activation, thereby suppressing MyD88-dependent TLR signaling (Xu et al., 2012). Moreover, MHC class I engagement can modulate type I interferon responses by recruiting SHP-2 to inhibit STAT1 phosphorylation, thus restraining antiviral signaling (Xia et al., 2019).

Our findings support that, similar to previous studies, Src might be involved in the phosphorylation of the tyrosine residue found in the cytoplasmic domain of MHC class I. Earlier work showed that the Rous sarcoma virus kinase pp60^v-src (rous sarcoma virus gene designated Src that encodes a 60 kDa phosphoprotein) can phosphorylate the conserved tyrosine residue in the MHC class I cytosolic tail *in vitro* (Guild et al., 1983). A Src-family kinase inhibitor was able to block this modification, indicating that Src kinases mediate the phosphorylation (Xu et al., 2012). Moreover, it was recently found that mutations in the cytosolic tyrosine alter CD8□ T cell responses. Using non-phosphorylatable or phosphomimetic tyrosine mutants of MHC class I expressed in APCs (human macrophage-derived cell line KG-1 and mouse DC line DC2.4), it was shown that phosphorylation of the cytosolic tyrosine markedly boosted CD8^+^ T cell priming, activation, and memory. In essence, this tyrosine not only governs MHC class I trafficking inside APCs, but also alters and reshapes antigen presentation and T cell activation (Sun et al., 2025). Collectively, these findings position reverse MHC class I signaling as a critical mechanism balancing cytoskeletal polarization required for specific antigen presentation with the need to prevent excessive inflammatory activation.

Our findings suggest that the molecular pathways regulating MTOC polarization are remarkably conserved among different immune cell types. Key components previously identified in T cells, appear to play a similarly important role in dendritic cells (Muntjewerff et al., 2020). These findings support the idea that fundamental mechanisms controlling cytoskeletal polarization are broadly shared within the immune system. This discovery broadens the current understanding of the signaling networks underlying immune cell polarity and points to a previously unappreciated role for MHC class I molecules in the regulation of cytoskeletal organization and directional cellular responses.

Finally, our data indicate that during the formation of multiple simultaneous T cell contacts, a situation arising during T cell scanning in the secondary lymph nodes (Bousso & Robey, 2003), the MTOC and associated secretory machinery are preferentially oriented toward the synapse exhibiting the strongest reverse MHC class I signaling. We propose that this selective polarization constitutes a decision-making mechanism by which DCs prioritize one T cell interaction among several competing contacts. This “DCision” mechanism would ensure that the DC directs its immunological resources, including MTOC alignment, Golgi orientation, and cytokine secretion, toward the T cell providing the strongest activation cues, thereby maximizing the efficiency and specificity of T cell priming.

## Acknowledgements

G.v.d. Bogaart is supported by the European Research Council (ERC) under the European Union’s Horizon 2020 research innovation program (grant agreement 862137). NR received funding by a Long-Term Fellowship from the European Molecular Biology Organization (EMBO-LTF, ALTF 232-2016) and a Veni grant from the NWO Talent Scheme (016.Veni.171.097).

## Methods

### Murine cell culture

C57BL/6 wild-type (WT) and OT-I mice were purchased from Charles River and hosted at the Central Animal Laboratory (CDL) of the Radboud University Medical Center. The mice were kept in top-filter cages and received a standard diet. At the time of an experiment, the mice were sacrificed by cervical dislocation at an age of 8-12 weeks old. All experiments in this project were approved by an animal ethical committee with regard to the care and use of animals.

For the generation of BMDCs, murine femurs and tibias were acquired from female WT mice. From the femurs and tibias, bone marrow derived stem cells were obtained and cultured in RPMI-1640 medium with 10% fetal bovine serum (FBS, 758093, Greiner bio-one), 1% antibiotic-antimycotic (15240-062, Gibco), 1% ultraglutamine (BE17-605E/U1, Lonza Bioscience), 50 μM β-mercaptoethanol (60-24-2, Sigma-Aldrich) and 200 ng/ml human Flt3L (130-096-479, Miltenyi Biotec) for eight days at 37ºC in an incubator with 10% CO_2_. After eight days, floating cells were collected and a CD45R (B220) depletion (microbead isolation kit (130-049-501), Miltenyi Biotec) was performed to remove plasmacytoid DCs from the cell population. For the isolation of naive CD8^+^ T lymphocytes, spleen and lymph nodes were isolated from female OT-I mice. The organs were homogenized and digested for 30 minutes at 37ºC by using a mixture of collagenase and DNAse, at 1 mg/ml and 130 μg/ml, respectively. CD8^+^ T cells were obtained after LD column magnetic antibody cell sorting (MACS, 130-042-901, Miltenyi Biotec) with anti-CD8 antibody beads from Miltenyi Biotec (130-104-075). The beads were used according to the company’s protocol.

### Human cell culture

Experiments were performed using monocyte-derived DCs and peripheral blood lymphocytes PBLs. Approval to conduct experiments with human blood samples was obtained from the Dutch blood bank, and all experiments were conducted according to national and institutional guidelines (NVT0459.00). Informed consent was obtained from all blood donors by the Dutch blood bank. Samples were anonymized and none of the investigators could ascertain the identity of the blood donors. Briefly, peripheral-blood mononuclear cells (PBMCs) were isolated from buffy coats by density gradient centrifugation. PBLs were separated from monocytes after positive selection of CD14^+^ cells with CD14 microbeads, LS Column and a MidiMACS Separator (Miltenyi Biotec Order no. 130-050-201). Monocytes were differentiated into moDCs by culturing in RPMI1640 medium supplemented with L-glutamine (21875-034; Gibco), 10% fetal bovine serum (FBS; 10309433; Thermo Fisher Scientific), 1% antibiotic antimycotic (AA; 15240062; Gibco), 300 U/ml interleukin-4 (130-093-924; Miltenyi), and 450 U/ml granulocyte macrophage colony-stimulating factor (130-093-867; Miltenyi) for 6 days at 37°C and 5% CO_2_.

### Mixed leukocyte reaction (MLR)

MLRs were performed by coculturing moDCs with PBLs. For the allogenic MLR reactions, human PBLs were obtained from different donors than the moDCs. The MLR assays were carried out on 12 mm diameter glass coverslips in 24-well plates for 2 hours to ensure efficient moDC-T cell contact in a 1:10 moDC to PBL ratio. Cells were cultured in RPMI1640 medium supplemented with L-glutamine, 10% fetal bovine serum, 1% antibiotic antimycotic in a total volume of 500 μl per well. For microtubule, kinase inhibition experiments were performed by adding nocodazole (30 µM) (Sigma Aldrich, Catalog. number M1404), Colchicine (50 µg/ml) (Sigma Aldrich C9754, CAS Number:64-86-8). MoDCs were preincubated for 1h with the inhibitors prior to co-incubation with PBLs for an extra one hour. For PBL treatment with IL-12, PBLs were treated with IL-12 (Immunotools, Cat.number 11349122) for one hour.

### BMDC-T cell synapse formation

Fresh day 8 Flt3L BMDCs (B220^-^) were activated in a non-adherent well plate (7007, Corning) for four hours at 37ºC and 5% CO_2_ with LPS (L4391-1MG, Sigma-Aldrich) and CpG ODN1826 (tlrl-1826-1, InvivoGen) both at a concentration of 1 μg/ml. The cells either received no addition of protein/peptide, OVA albumin (0.5 mg/ml, 321001, Lionex GmbH), OVA in combination with IgG against OVA (20 μg/ml OVA + 500 μg/ml Goat polyclonal IgG against chicken ovalbumin, 0855303, MP Biomedicals) or SIINFEKL (500 ng/ml, OVA_257-264_, AS-60193-5, Tebu-Bio). OVA and OVA-immunocomplexes were added for the full activation time, while SIINFEKL was added for the last 30 minutes of activation. After activation, OT-I T cells obtained from mouse spleen and lymph nodes and labelled with CFSE (C34554, Invitrogen) were added 2:1 to the BMDCs and placed on PLL-coated coverslips. The DCs and T cells were incubated for 1 hour at 37°C and 5% CO_2_. After this time the cells were fixed for 20 minutes with 4% PFA.

### BMDC-bead synapse formation

Fresh day 8 Flt3L BMDCs (B220^-^) were activated in a non-adherent well plate for four hours at 37ºC and 5% CO_2_ with TLR4 agonist LPS and TLR9 agonist CpG-B ODN1826 both at a concentration of 1 μg/ml. During activation, 6 μm streptavidine beads were coated with different biotinylated antibodies (3.75 μg of antibody per 6 million beads, see table 1.) and incubated for 30 minutes in PBA (PBS + 1% BSA and 0.05% sodium azide) at room temperature (RT). After activation, the BMDCs were mixed with beads on poly-L-lysine (PLL, 25988-63-0, Sigma-Aldrich) coated coverslips in a 1:1 ratio and were incubated for 30 minutes at 37ºC and 5% CO_2_. After this time the cells were fixed for 20 minutes with 4% paraformaldehyde (PFA) and stained.

**Table 1.**
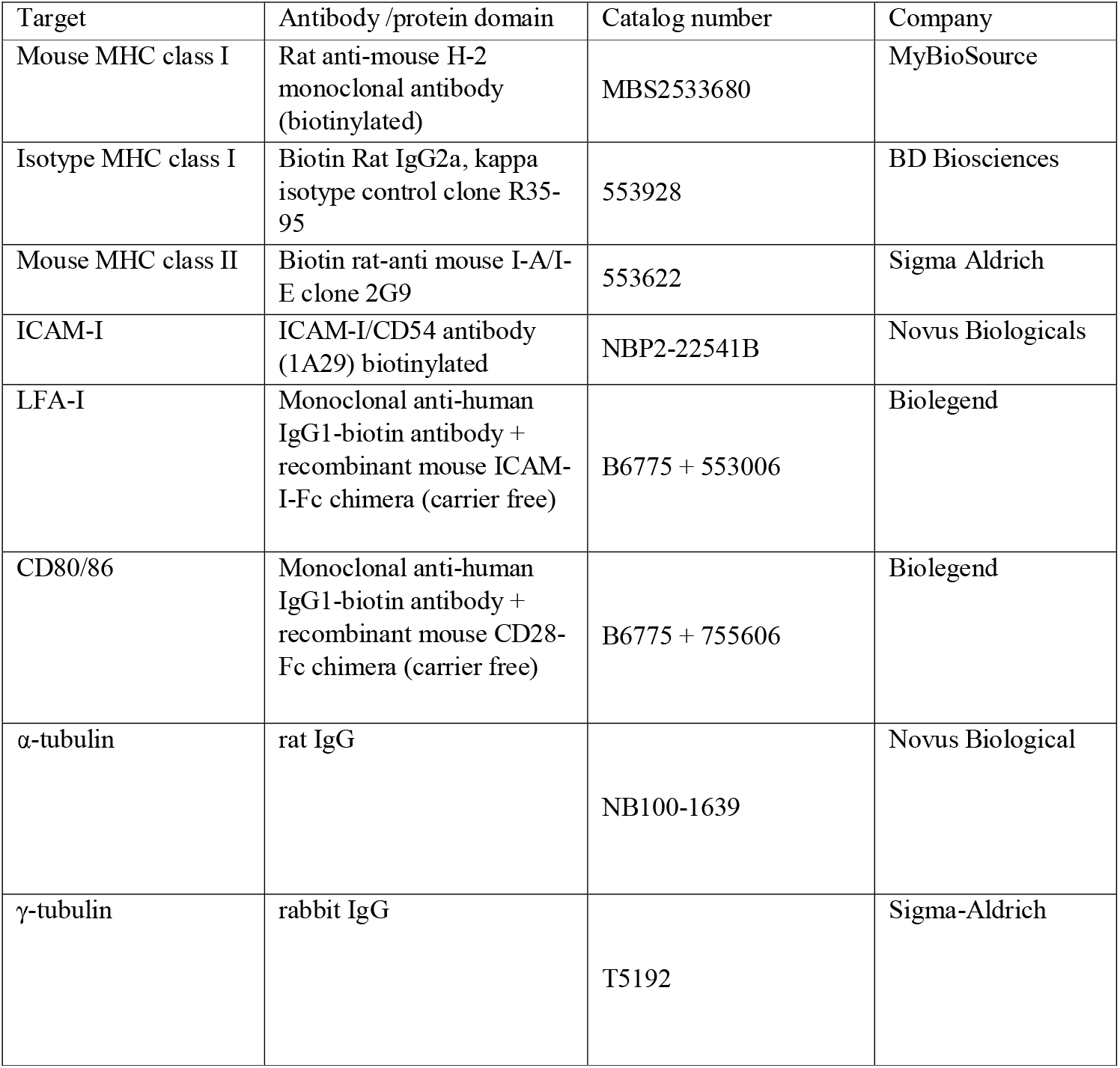
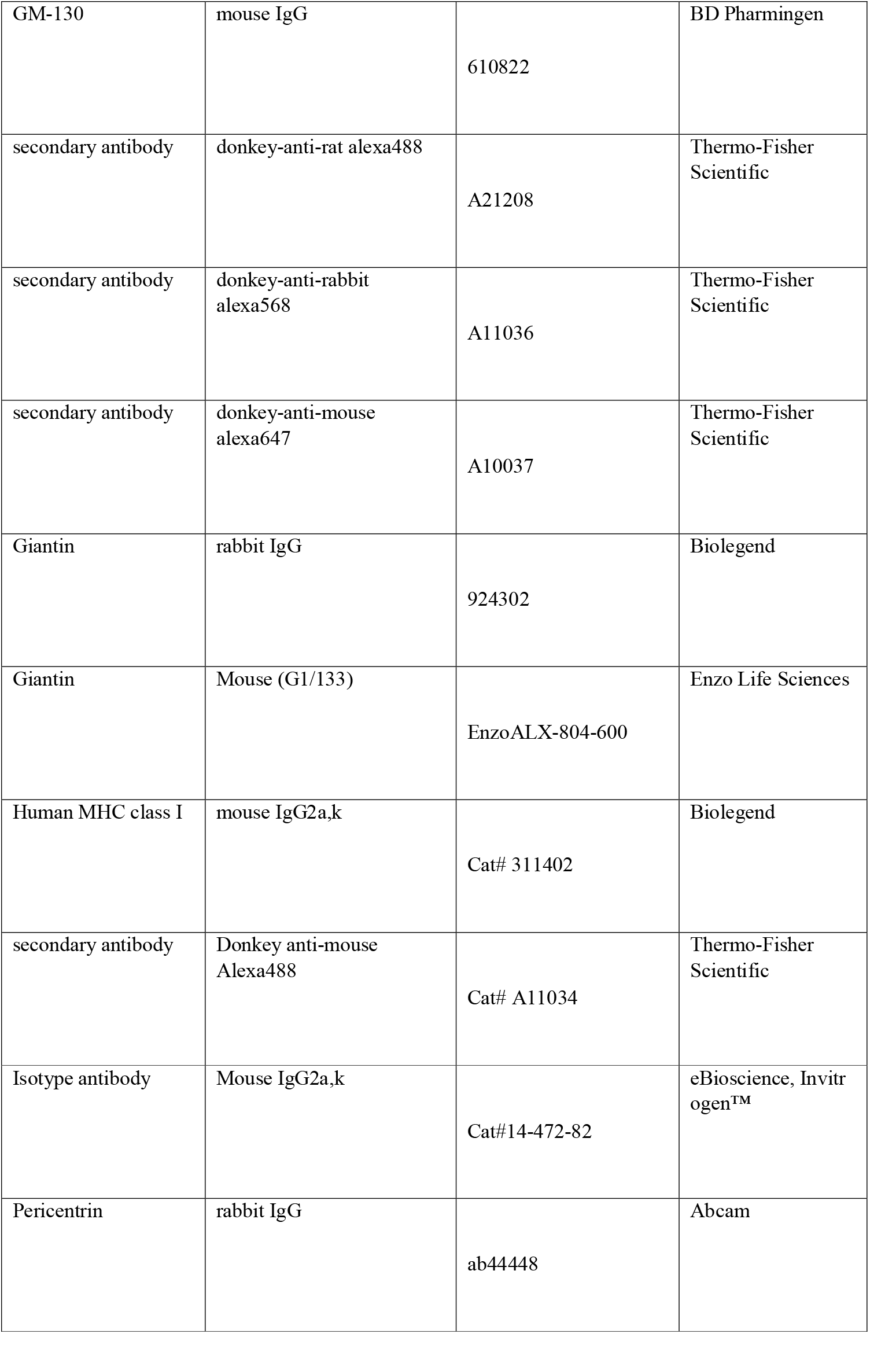
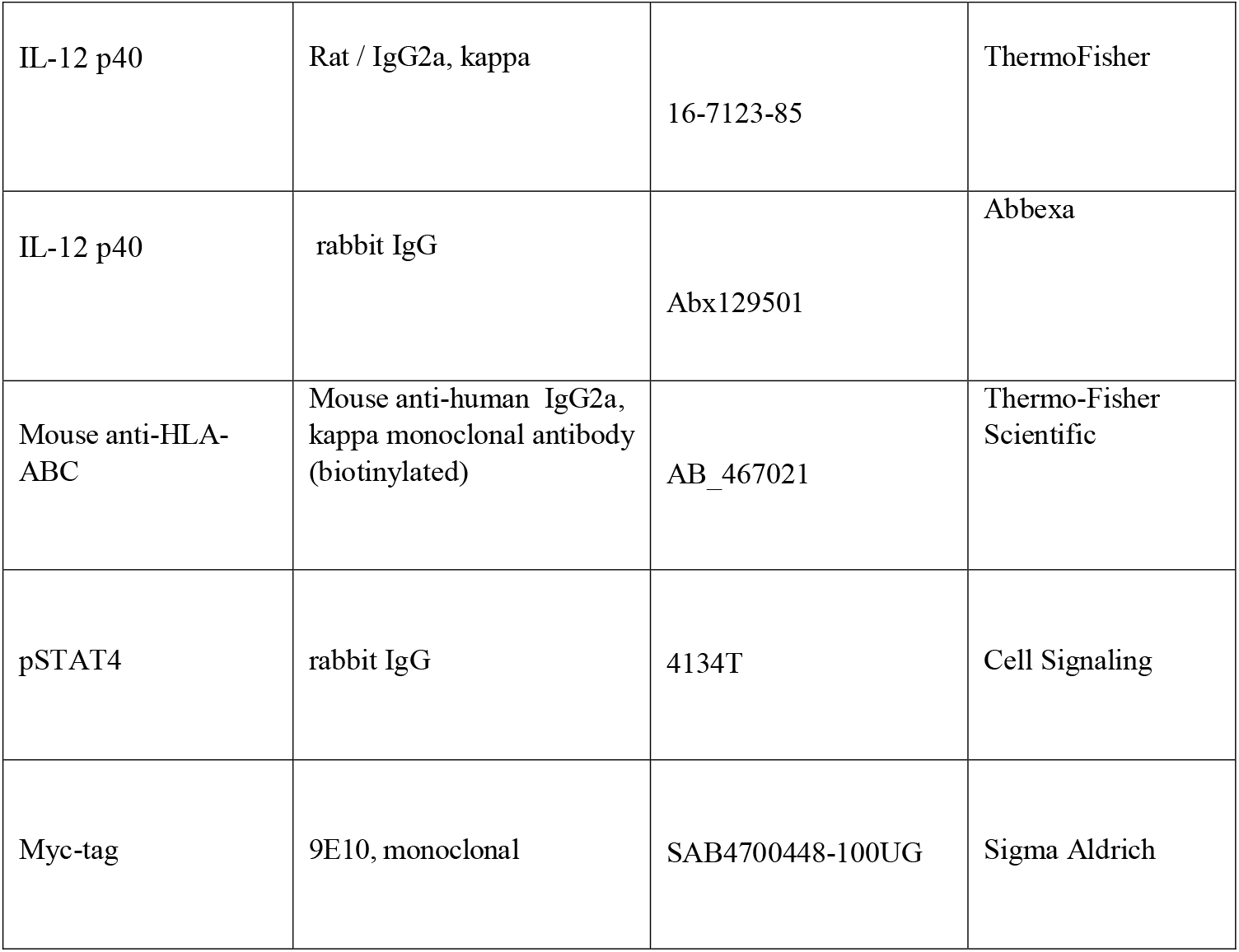
The antibodies and protein domains used in the experiments.

### MoDC-bead synapse formation

Day 6 moDCs were activated in a non-adherent well plate for four hours at 37ºC and 5% CO_2_ with LPS and TLR7/8 agonist R848 at a concentration of 1 μg/ml and 2.5 mg/ml respectively. After activation, the moDCs were mixed with beads coated with either an antibody against MHC class I (13-9983-82, ThermoFisher) or its corresponding isotype control (13-4724-85, ThermoFisher) on PLL-coated coverslips in a 1:1 ratio and were incubated for 30 minutes at 37ºC and 5% CO_2_. After this time the cells were fixed for 20 minutes with 4% PFA.

### Inhibitors-treatments

After 2 hours incubation with nocodazole (5 µM) (Sigma Aldrich, Catalog. number M1404), Colchicine (50 µg/ml) (Sigma Aldrich C9754, CAS Number:64-86-8), Src I1 Dual site Src kinase inhibitor (5 μM) (Bio-Techne, Catalog. number 3642), PP1 (5uM) (Bio-Techne Catalog. Number 139), phospholipase C inhibitor (U73122) (0.4 uM) (Bio-Techne Catalog. Number 1268) and its negative control compound (U73343) (0.4 uM) (Bio-Techne Catalog. Number 4133), the PKC inhibitor Gö6983 (Bio-Techne Catalog. Number 2285), the beads coated with antibody against MHC class I or naked beads were added in BMDCs/moDCs and MTOC polarization was assessed using immunofluorescence microscopy.

### Immunofluorescence staining

After fixation with 4% PFA in PBS for 20 minutes at RT, excess PFA was quenched with a solution containing 100 mM glycine and 100 mM NH_4_Cl in PBS for 20 minutes at RT. Human DCs were blocked-permeabilized with 2.5% BSA + 1% donkey serum (017-000-121-, Jackson) + 0.15% Triton X-100 in PBS. Mouse DCs were blocked-permeabilized with 2.5% BSA + 1% donkey serum + 0.1% Triton X-100 in PBS. Synapses from murine BMDCs were stained with antibodies against α-tubulin (1:500, rat IgG, NB100-1639, Novus Biological), γ-tubulin (1:500, rabbit IgG, T5192, Sigma-Aldrich) and the cis-Golgi marker GM-130 (1:100, mouse IgG, 610822, BD Pharmigen) for one hour at RT. Human cells were stained for α-tubulin (1:500, rat IgG, NB100-1639, Novus Biological), γ-tubulin (1:500, rabbit IgG, T5192, Sigma-Aldrich), pericentrin (1:100, rabbit IgG, ab44448, Abcam) and the Golgi marker Giantin (1:100, mouse (G1/133), EnzoALX-804-600 and 1:100, rabbit IgG, Biolegend 924302) for one hour at RT. After washing, the synapses were stained with fluorescently labelled secondary antibodies: donkey-anti-rat alexa488 (1:400, A21208, Thermo-Fisher Scientific), donkey-anti-rabbit alexa568 (1:400, A11036, Thermo-Fisher Scientific) and donkey-anti-mouse alexa647 (1:400, A10037, Thermo-Fisher Scientific). After incubation with the secondary antibody, the cells were washed with PBS and mounted with a glycerol-containing mounting medium (68 g glycerol, 100 mM NaPi pH 7.4 + trollox + DAPI).

### Transfections of RAW 264.7 cells

RAW 264.7 cells were transfected by electroporation using the NEON Transfection System (Invitrogen) with 10 μl NeonTips (MPK1025; Thermo Fisher Scientific). The protocol entailed 1 pulse of 1,680 V and 20 ms. The concentration of DNA used was 3 μg per 1 million cells. The pCEP4-myc-HLA-A2 construct was obtained from Addgene (#135504) (McShan et al., 2019). The pCEP4-myc-HLA A2_Y344F construct was generated by point mutagenesis and has been deposited at Addgene. The cells were seeded and mixed with beads on poly-L-lysine (PLL, 25988-63-0, Sigma-Aldrich) coated coverslips in a 1:1 ratio and were incubated for 30 minutes at 37ºC and 5% CO_2_. After this time the cells were fixed for 20 minutes with 4% paraformaldehyde (PFA) and immunostained as described above.

### Confocal microscopy

Synapses were imaged on an SP8 confocal microscope with a 63 × 1.2 NA water immersion objective (Leica) or Zeiss LSM 800 microscope.

### Transmission electron microscopy

Sapphire discs were incubated with polyethylamine (0.2% PEI) in H_2_O for 1 min. Discs were placed in a 24-well dish, rinsed twice with H_2_O, and dried at RT. MoDCs from donor 1 together with PBLs from donor 2 were seeded in RPMI complete medium and allowed to interact for 2 h. Samples were immediately transferred to a LEICA EM ICE machine for high-pressure freezing. Freeze substitution was performed in a Leica automated freeze substitution machine according to the following protocol: 33 h at −90°C with 4% uranyl acetate in methanol and osmium tetroxide the temperature was then raised to −20°C, held for 14 h, and then raised to 5°C. Embedding was started with acetone washes and infiltration of the samples with Epon 812 substitute like Glycid Ether, with hardener (MNA) and accelarator (DMP-30) mixtures with increasing amounts of EPON. Samples were infiltrated with pure EPON overnight and then allowed to polymerize at 60°C for 2–3 d. Serial sections of 70 nm were cut with an automated tape collecting ultra-microtome (Leica, UC6). Ultrathin sections were contrasted with uranyl acetate and lead citrate and examined under the ThermoFisher Talos L120C Transmission Electron Microscope.

### Microcontact printing

Sylgard 184 (SYLGARD 184, Sigma Aldrich, 761028-5EA), containing 10 parts of elastomer with 1 part current agent, was poured in a master mold for producing a stamp with pillars with a period of 16 µm and a diameter of 8 µm (ThunderNIL). The mold was placed in a vacuum system for 30 min at 80 °C and then was cured overnight at 80 °C. The stamp was placed with the pillars facing upwards and was covered with MHC class I (Biolegend, 311402) or Isotype antibody (Invitrogen™, 14472482) at a concentration of 100 μg/mL together with the 10 μg/mL Donkey anti-mouse Alexa488 (Thermo-Fisher, A11034) in PBS for 1 h in the dark to avoid photobleaching of the fluorescent label. Then the stamps were briefly cleaned with water and dried under a N_2_ steam. Cleaned coverslips were placed on top of the stamps facing the motif. Each coverslip was lightly pressed with tweezers onto the stamp for approximately 1 minute. The stamp was removed from the coverslips and the cells were immediately loaded (10^5^ cells per coverslip) onto the coverslip in a 24-well plate. Cells were incubated for 30 or 60 min and then fixed with 4% PFA diluted in PBS and immunostained for pericentrin, as described above.

### Statistical analysis

The plots for polarization index show the average ± standard error of the mean (SEM). Normality was checked by D’Agostino & Pearson normality test, followed by either a Student t test or a one-way analysis of variance (ANOVA) followed by Dunnet multiple comparisons, as specified in the figure legends. The significance level was set at 5% (p = 0.05).

## Figure legends

**Supplementary Figure 1.**
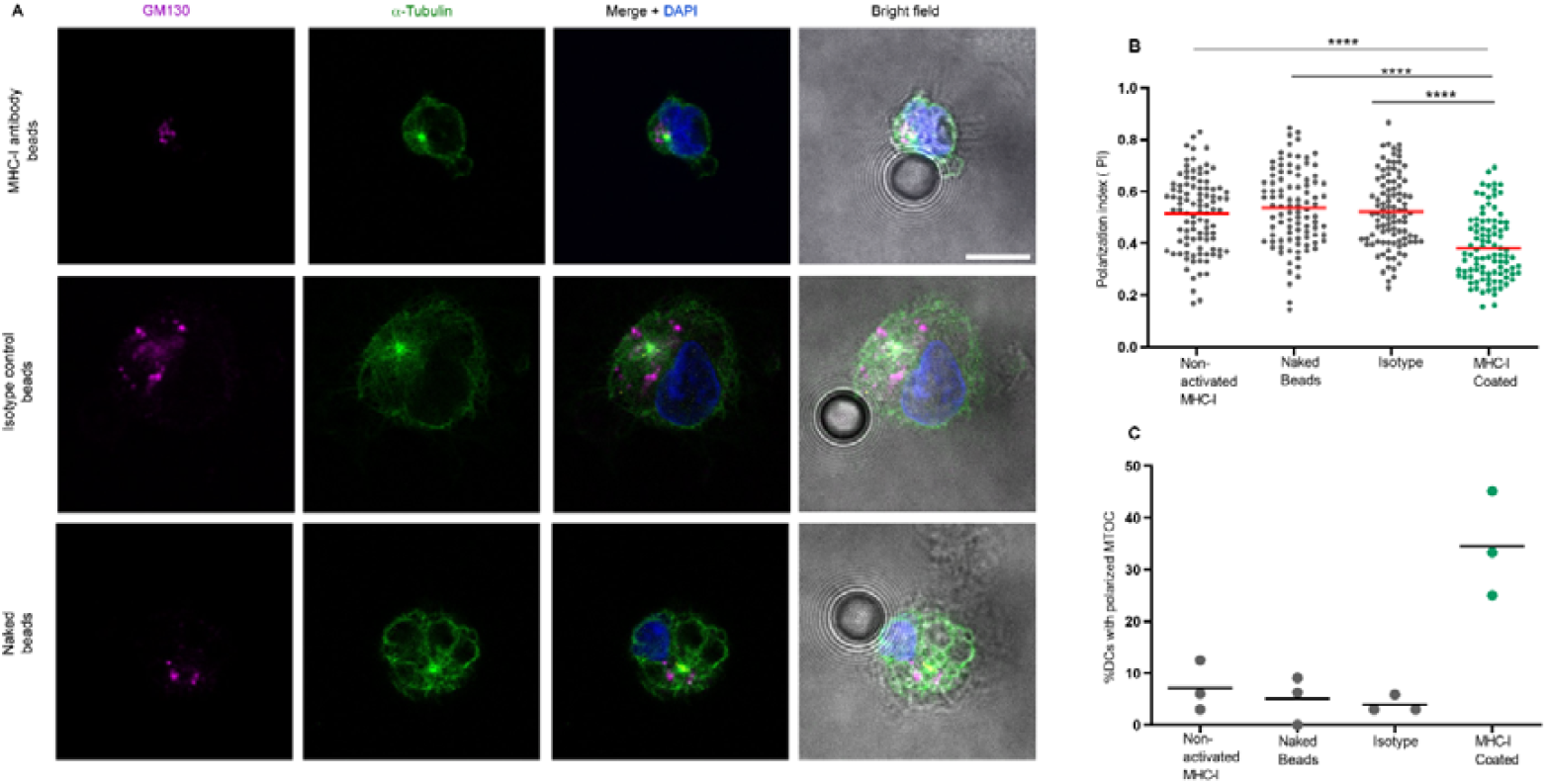
Reverse MHC class I signaling drives MTOC polarisation in human DCs. **(A)** Representative confocal microscopy images of the human monocyte-derived DCs (moDCs) incubated with naked beads, beats coated with isotype antibody, or beads coated with antibodies against MHC class I. Cells were immunostained for cis-Golgi marker GM-130 (magenta) and α-tubulin (green). Scale bar, 10 µm. **(B)** Analysis of MTOC polarization in moDCs using beads coated with isotype or antibody against MHC class I. One-way ANOVA with Dunnett’s multiple comparison, Mean ± standard error of the mean ****P<0.0001). **(C)** Same as panel B but now displayed as percentage polarized cells (defined as polarisation index < 0.3) for individual donors.

**Supplementary Figure 2.**
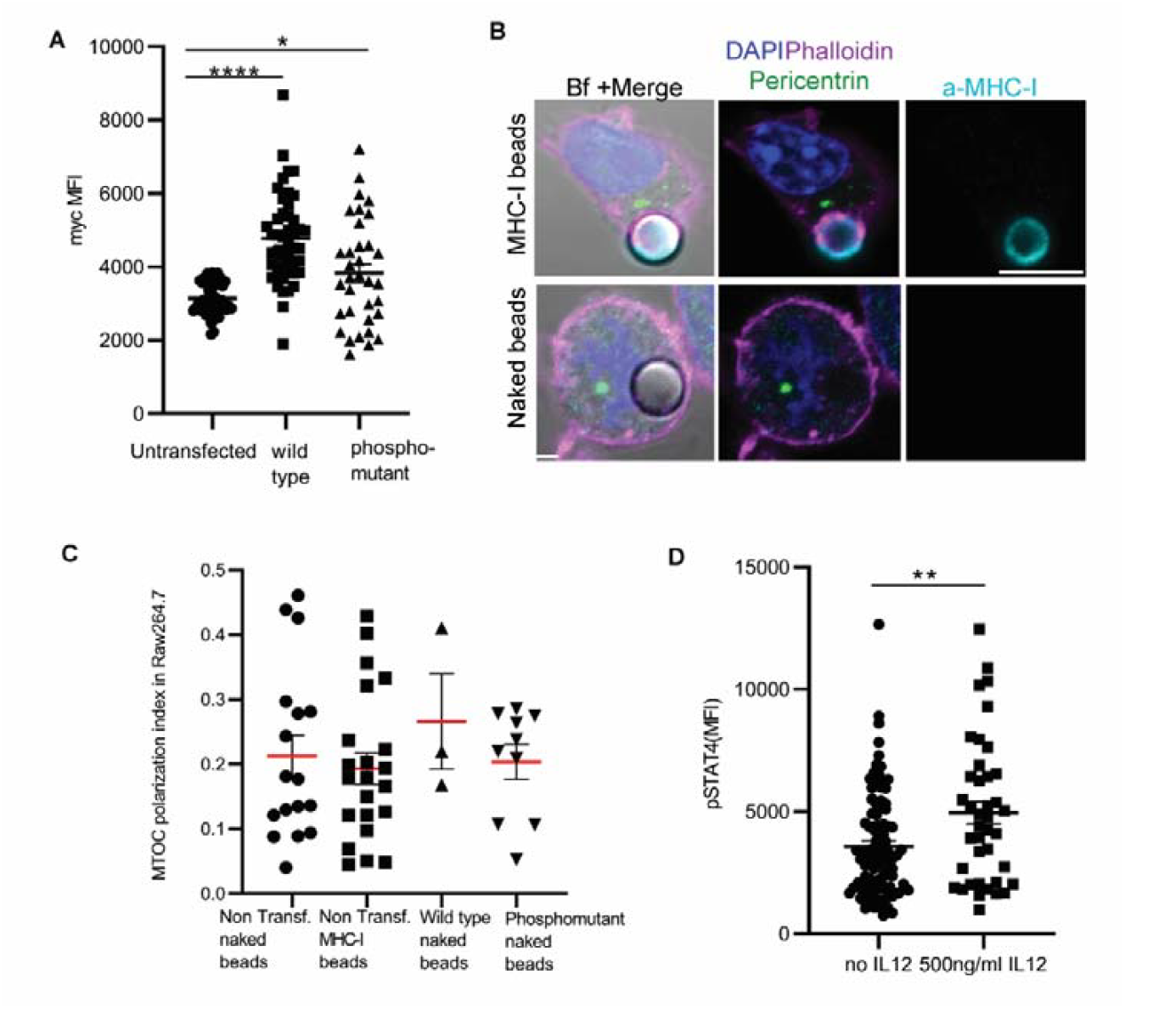
Reverse MHC class I signaling drives MTOC polarisation in human DCs. **(A)** Quantification of Myc mean fluorescence intensity (MFI) in untransfected or heterologously expressing human HLA-A*02:01 Raw 264.7 macrophages. Unpaired t test; ****, P < 0.0001. **(B)** Representative confocal microscopy images of the RAW 264.7 macrophages heterologously expressing human HLA-A*02:01 and incubated with bead coated with antibodies against human (but not mouse) MHC class I or with naked beads. Cells were immunostained for DAPI (blue), pericentrin (green), MHC class I (cyan) and phalloidin (magenta). Scale bars, 10 µm. **(C)** Quantification of the MTOC polarization in non-transfected, wild type, phosphodead Raw 264.7 macrophages incubated with beads coated with antibodies against MHC class I or naked beads. **(D)** Quantification of pSTAT4 MFI in PBLs incubated with human IL-12. Unpaired t test; **, P < 0.01.

